# Metabolic Segregation and Functional Gene Clusters in Anaerobic Digestion Consortia

**DOI:** 10.1101/2024.08.04.606487

**Authors:** Yubo Wang, Ruoqun Zhang, Chunxiao Wang, Weifu Yan, Tong Zhang, Feng Ju

**Affiliations:** Environmental Microbiome and Biotechnology Lab, School of Engineering, Westlake University, Hangzhou 310024, Zhejiang Province, China; Environmental Microbiome Engineering and Biotechnology Laboratory, The University of Hong Kong, Pokfulam, Hong Kong; Westlake Laboratory of life Sciences and Biomedicine, School of Life Sciences, Westlake University, Hangzhou 310024, China

**Keywords:** Metabolic segregation, H_2_ and formate, Functional gene clusters, Genome-centric transcriptome, Enriched microbiota, Anaerobic digestion

## Abstract

Enrichment experiment integrated with genome-centric meta-omics analysis revealed that metabolic specificity, rather than metabolic flexibility, governs the anaerobic digestion (AD) ecosystem. This study provides new insights into interspecies electron transfer in the AD process, highlighting a segregation in the metabolism of H₂ and formate. Our findings indicate that H₂ serves as the primary electron sink for recycling redox cofactors, NAD⁺ and oxidized ferredoxin (Fd_ox_), during primary fermentation. In contrast, formate predominates in the co-occurring secondary fermentation, particularly under conditions of elevated H₂ concentrations. Notably, no evidence of biochemical interconversion between H₂ and formate was identified in the primary fermenting bacteria or in syntrophs enriched in this study. This segregation of H₂ and formate metabolism likely benefit the anaerobic oxidation of butyrate and propionate with a higher tolerance to H₂ accumulation. Furthermore, this study underscores the functional partitioning among microbial populations within key niches of the AD bioprocess: primary fermentation, secondary fermentation (syntrophic acetogenesis), hydrogenotrophic methanogenesis, and acetoclastic methanogenesis. Through genome-centric analysis of the AD microbiome, we identified several key functional gene clusters, highlighting the potential for improving genome-centric genotype-phenotype correlations, which is particularly valuable for strict anaerobes that remain challenging to isolate and characterize in pure culture.

## 1. Introduction

Anaerobic digestion (AD) has been of central importance in the field of environmental biotechnology since 1980s, being applied to stabilize organic matter in high-strength waste streams, to reduce biosolids and support bioenergy production [1–6]. Despite being a relatively mature engineering technology, challenges persist, particularly process upsets such as over-acidification and foam formation [7], both of which can be induced by the accumulation of volatile fatty acids (VFAs) arising from an imbalance between the rapid acidogenesis and the slower β-oxidation. Fine-tuned control and optimization of the AD process depend on a deeper understanding of the sequential microbial reactions mediated by anaerobic populations across the four function niches: hydrolysis (conversion of polymers to monomers), primary fermentation/acidogenesis (conversion of monomers to H_2_/HCOOH, acetate and other reduced fatty acids), secondary fermentation (conversion of reduced fatty acids to acetate and H_2_/HCOOH) and methanogenesis (CH_4_ generation mainly from acetate and H_2_/HCOOH) [8–14].

Among the four function guilds, anaerobes mediating secondary fermentation has gained significant scientific attention due to their intriguing bioenergetic strategies and ability to thrive under thermodynamic boundary conditions [15, 16] [17, 18]. Secondary fermentation is often regarded as the bottleneck step in the AD bioprocess [19], yet largely unknown is the phylogenetic and metabolic diversity of anaerobes in this function niche. How prevalent is obligate syntroph compared to syntrophy in opportunistic microbes [17, 20, 21]? Many uncharacterized and uncultivated bacteria in the AD consortia have yet-to-be-identified relationships with the two trophic groups, i.e., primary fermentation *vs.* secondary fermentation, and an in-depth understanding on the metabolic flexibility or specificity of the syntrophic populations could help elucidate the intricate ecology that supports the methanogenic conversion of organic compounds.

One of the objectives of this study is to chart the metabolic flexibility of syntrophs, focusing on <u>s</u>yntrophic <u>b</u>utyrate <u>o</u>xidizing <u>b</u>acteria (SBOBs) and <u>s</u>yntrophic <u>p</u>ropionate <u>o</u>xidizing <u>b</u>acteria (SPOBs). As key intermediates in the anaerobic mineralization of organic compounds, propionate and butyrate can account for 20-43% of the total CH_4_ formation [22]. We aim to assess the flexibility of SBOBs and SPOBs in switching their substrate spectrum from reduced VFAs (e.g., butyrate, propionate) to fermentative substrates (e.g., crotonate, fumarate, glucose). Do they prioritize fermentative substrates—due to their higher energetic potential and biomass yield—or continue to utilize VFAs even when fermentative substrates are available?

Another well-accepted metabolic trait of the SBOBs and the SPOBs is their sensitivity to product inhibition by H_2_. H_2_ concentration as low as 100 Pa (∼1000 ppm) can inhibit the anaerobic oxidation of propionate or butyrate[22]. In the established scheme of anaerobic carbon cycling, almost all substrates, including carbohydrates and amino acids, generates H_2_ as a byproduct [23, 24]. How does such H_2_ production affect the cooccurring H_2_-sensitive oxidation of butyrate and propionate? Systematic characterization of H_2_ dynamics (generation and turnover) during AD remains limited, with most studies focusing on fermentation without methanogenesis [25]. The impact of H₂ accumulation on anaerobic VFA oxidation in these ecosystems has been insufficiently evaluated [17, 26–28]. Interestingly, experimentally determined H₂ inhibition concentrations—2.03 kPa for butyrate oxidation and an H₂ inhibition constant (Kₚ(H₂)) of 0.11 atm (11.1 kPa) for propionate oxidation—are far higher than the theoretical threshold of 100 Pa [25, 29, 30]. Additionally, emerging evidence suggests that formate, rather than H₂, may play a more significant role as an interspecies electron carrier in the syntrophic oxidation of organic acids and alcohols, such as propionate and 1,3-propanediol [31, 32].So, the second objective of this study is to address this knowledge gap by integrating the characterization of H_2_ dynamics with the degradation dynamics of VFAs in lab-scale AD reactors, and to evaluate the relevance of H_2_ and formate as interspecies electron carriers through the transcription profile of formate dehydrogenase (FDH) and hydrogenase (H_2_ase).

To facilitate characterization, we began by enriching microbial consortia dominated by primary fermenting bacteria, SBOBs, and SPOBs. This enrichment was conducted using three sets of reactors, each fed with a different substrate: glucose, butyrate, and propionate, respectively. We then evaluated the metabolic versatility of the enriched anaerobes by examining their metabolic capacity and transcriptomic response to fermentative substrates (i.e., glucose, fumarate, crotonate) and VFAs (i.e., propionate and butyrate), respectively. Throughout the batch experiments, we monitored the dynamics of H₂ generation and H₂ turnover as an intermediate metabolite, as well as whether the syntrophic oxidation of propionate and butyrate can proceed under high concentration of H₂ accumulation.

## 2. Material and methods

### 2.1 Enrichment experiments

Three sets of AD consortia were enriched for the efficient methanogenic conversion of glucose, propionate and butyrate, respectively. Duplicate enrichments utilizing each substrate were processed in 1000-ml serum bottles with working volume of 500 ml. Hereafter, the consortia enriched with glucose as substrate will be referred to as ‘Consortia-G’, the consortia enriched with butyrate as substrate will be referred to as ‘Consortia-B’, and the consortia enriched with propionate as substrate will be referred to as ‘Consortia-P’.

Trace elements and mineral nutrients were prepared following the DSMZ medium recipe119 for *Methanobacterium*^1^, with one slight modification: 0.4 g/L MgSO₄·7H₂O was replaced by 0.33 g/L MgCl₂·6H₂O to maintain the sufficient supply of Mg^2+^ while at the same time minimize the potential for sulfate reduction within the consortia.

The inoculum used in this enrichment experiment was surplus sludge sampled from Chengxi wastewater treatment plant (Hangzhou, China). Inoculum biomass pellet was obtained from 500 ml surplus sludge after centrifugation at 1000 rpm for 10 min, this biomass pellet was resuspended in solution medium plus trace element solution to 500 ml and served as inoculum. Serum bottles in which the enrichment experiments happened were initially sparged with N_2_ gas to maintain an anaerobic environment.

Over the 219-day acclimation and enrichment period, the organic loading rate for each substrate was progressively increased from 62 mg-COD/g-VS/d to 259 mg- COD/g-VS/d (Figure S1). The substrate was supplemented every other day in a fill-and-draw mode, i.e., 1 ml or 2 ml of the concentrated (180 g-COD/L or 360 g-COD/L) solution of each substrate would be diluted into 20 ml nutrient solution and be supplied into the reactor to replace an equal volume, that is 20 ml of the supernatant after sludge settling. In this way, nutrient solution will be refreshed every 50 days. No biomass was intentionally wasted during the reactor operation, and a rough estimation of the SRT at ∼50 days was achieved by biomass loss during sampling for routine VS/TS measurement and for DNA archive. No stirring was introduced in this experiment and the serum bottles were shaken continuously at 150 rpm in a thermostat water bath and the temperature was maintained at (35±1) °C.

### 2.2 Batch test on the metabolic flexibility of the primary fermenting bacteria, SBOBs and SPOBs enriched

Following enrichment, an experiment was conducted to characterize the metabolic flexibility of anaerobes within three functional guilds: primary fermentation, syntrophic butyrate oxidation, and syntrophic propionate oxidation. The experimental design is schematically illustrated in Figure S2. Specifically, five batch tests were performed, with five different substrates sequentially applied to each of the three enriched consortia. For each batch test, eight serum bottles were used, with each bottle corresponding to a specific sampling time point. At each time point, 30 mL (2*15ml) of sludge was collected and flash frozen for total RNA extraction.

Take the batch test conducted on SBOBs for example, 1000 mL of enriched sludge (Consortia-B) was obtained after 219 days by combining 500 mL from each of the two parallel reactors where SBOBs were enriched. The combined sludge was evenly distributed across eight serum bottles, each with a working volume of 125 mL. In the five sequential batch tests, the primary substrate for Consortia-B was switched from *n*-butyrate to propionate, then to glucose, crotonate, and finally to a tri-substrate mix of glucose, propionate, and *n*-butyrate (COD ratio 1:1:1). At the specified time points summarized in Figure S3, S4, S5, biogas composition in the gaseous phase and VFA profiles in the liquid phase were analysed. Based on preliminary results, such as butyrate degradation occurring under high H₂ concentrations, total RNA extraction was conducted only at the time points indicated by purple arrows in Figure S3, S4, S5.

### 2.3 Biomass archive, genomic DNA and genomic RNA extraction

To profile the succession of the microbial community during the enrichment process, 1.5 mL of sludge was archived weekly or biweekly from each reactor for sequencing-based analysis. In total, 115 sludge samples (19 time points × 6 reactors + 1 seed sludge) were archived for total genomic DNA extraction using the FastDNA SPIN Kit for Soil (MPBio, Santa Ana, CA, USA). DNA extracted from sludge collected at the same time point from two parallel reactors was pooled.

All genomic DNA extracts were subjected to polymerase chain reaction (PCR) amplification of the V4–V5 region of the universal 16S rRNA genes using the 515F-Y/926R primer set [33], followed by amplicon sequencing via the Novaseq-PE250 sequencing platform. Additionally, genomic DNA extracted from the seed sludge (day 0) and from sludge collected on days 85, 134, and 219 of the enriched consortia were subjected to metagenome sequencing.

Metatranscriptome analysis is relevant only in the succeeding batch tests. Sludge samples (15 ml *2) collected were flash frozen in liquid nitrogen and stored in -80 °C freezer. Total RNA extraction, sequencing and metatranscriptome analysis was conducted on twenty of the sludge samples collected. Details on the physiochemical parameters measured at each of the twenty time points have been summarized in Table S1 and Figure S3, S4, S5.

MagAttract PowerMicrobiome RNA kit (Qiagen, USA) was used for total RNA extraction, per the manufacturer’s instructions. Both the DNA extracts and the RNA extracts were quality-checked using agarose gel electrophoresis. Fragment sizing of the total RNA extracts was further checked using Agilent Bioanalyzer 2100 (Santa Clara, CA, USA), and the RNA integrity numbers (RIN) measured were in the range of 4.9-9.1 (Appendix file 1). Ribosomal rRNA was removed from the total RNA prior to cDNA generation, a process performed by technicians at Personal Gene Technology Co., Ltd. (Nanjing, China, all sequencing services involved in this study are provided by this company). Stranded DNA-seq and cDNA of the mRNA-seq were subjected to Illumina-PE150 platform to generate paired-end 150 bp reads, with the average insert size around 450 bp.

### 2.4 Meta-omics analysis

#### 2.4.1 16S rRNA amplicon sequence analyses

All amplicon sequences were processed using Qiime2 (Version 2021.4) [34]. Sequencing quality control was performed with DADA2 [35], and taxonomic assignment of the representative single gene variants (SVGs) was conducted using the Silva 138 SSU database. To assess how different substrates influence the enriched AD microbiomes, principal coordinate analysis (PCoA) was performed to visualize the dissimilarities among the three consortia enriched with glucose, propionate, and butyrate, respectively. In this PCoA, read counts were subsampled to 86,855, the minimum sequencing depth among the 57 samples after quality control. The PCoA analysis was based on Euclidean metrics with Hellinger distance.

#### 2.4.2 Genome-centric metagenome and metatranscriptome analyses

As summarized in Appendix file1, shotgun metagenome sequencing generated a total of 129 Gbp raw reads from 10 sludge samples, and 128 Gbp clean reads retained after quality-filtration. A detailed description on quality-control of the meta-omics data, assembly strategy, genome binning and genome dereplication could be found in the supporting information. Below is a summary of the key workflow and the bioinformatic tools used.

Ten single assembly and three co-assembly of the clean reads were through the metaWRAP-Assembly module megahit [36]. QUAST [37] evaluation on the contig sequences assembled could be found in Appendix file1. Genome binning was conducted with the metaWRAP binning module metabat2 [38]. The generated metagenome assembled genomes (MAGs) were dereplicated using the dRep tool [39]. Quality of MAGs was assessed using CheckM v1.0.7, based on the presence of lineage-specific, conserved, single-copy marker genes [40].

Taxonomic affiliation of the MAGs was inferred using GTDB-Tk v0.1.3 under ‘classification’ mode, with 120 ubiquitous bacterial single-copy protein sequences, and 122 ubiquitous archaeal single-copy protein sequences [41]. Genome annotation was performed using Prokka v1.12 [42], with the bacteria and archaea database for the annotation of bacterial MAGs and the archaeal MAGs, respectively. Open reading frame (ORF) calling was performed using Prodigal v2.6.2 [43]. Multiple databases were applied in the annotation of the ORFs, i.e., Uniprot database (release 2021-04), KOfam (release 2021-11). Putative hydrogenase genes were manually checked with HydDB classifier [44], and key functional genes with high transcription activity were also manually checked through on-line blast against the NCBI RefSeq non-redundant proteins database.

Abundance of the microbial population corresponding to each MAG was calculated by mapping clean DNA paired end reads to contigs of the MAGs assembled. And transcription activity of the function genes and of microbial population corresponding to each MAG was evaluated by mapping clean mRNA paired end reads to open reading frames (ORFs) within each MAG. The sequence Alignment and Map (SAM) formatted files generated by Bowtie 2 (v.2.4.3) [45] were converted to binary alignment and map (BAM) format and sorted using samtools (v.1.12) [46]. Methods for DNA-RPKM and RNA-RPKM calculation was the same as detailed in our previous omics-analysis work [47].

## 3. Results and discussion

### 3.1 Effects of glucose, butyrate and propionate on the succession of the anaerobic consortia enriched

Slow growth rates of the SBOBs and the SPOBs, and the selecting pressure imposed by SCFAs as the primary substrate is reflected by the significant decrease in the volatile suspended solids (VSS, a surrogate measure of the microbial biomass) in Consortia-B and Consortia-P. As summarized in Figure S6, at the same organic loading rate in terms of COD/L/d, in contrast to ∼43% increment in VSS from 8.0 g/L to 11.7 g/l in Consortia-G, both Consortia-B and Consortia-P are observed with a significant decrease in VSS by ∼50% from 8.0 g/l to 4.1∼4.4 g/l.

Figure 1 summarizes the CH₄ yield (270–350 mL/g-COD) and the CH₄ and H₂ contents in the biogas produced during the methanogenic conversion of glucose, butyrate, and propionate by the three enriched consortia on day 219. Interestingly, the methanogenic conversion of butyrate and propionate outperforms that of glucose, as evidenced by higher CH₄ content and superior CH₄ production rates and yields. While a direct comparison among these three substrates, each degraded by distinct consortia, may not be entirely equitable, the findings suggest that butyrate and propionate degradation is not necessarily slower than glucose degradation, despite their lower energy yield and weaker thermodynamic driving force. Beyond bioenergetics, the high abundance of functional populations, such as SBOBs in Consortia-B and SPOBs in Consortia-P, likely plays a pivotal role in driving the efficient anaerobic oxidation of butyrate and propionate by the acclimated consortia.

As expected, a distinct shift in microbiome composition (Figure 2) was observed during the enrichment process, with each substrate selectively enriching specific microbial populations. According to the PCoA analysis summarized in Figure 3, the genera characteristically enriched in Consortia-G include *Clostridium sensu stricto 3*, *Propionimicrobium*, and an unclassified genus within the family *Propionibacteriaceae* (hereafter referred to as *f_Propionibacteriaceae*). In contrast, the genus characteristically enriched in Consortia-P is *Smithella*, while Consortia-B is specifically enriched with *Syntrophomonas*.

**Figure 1.**
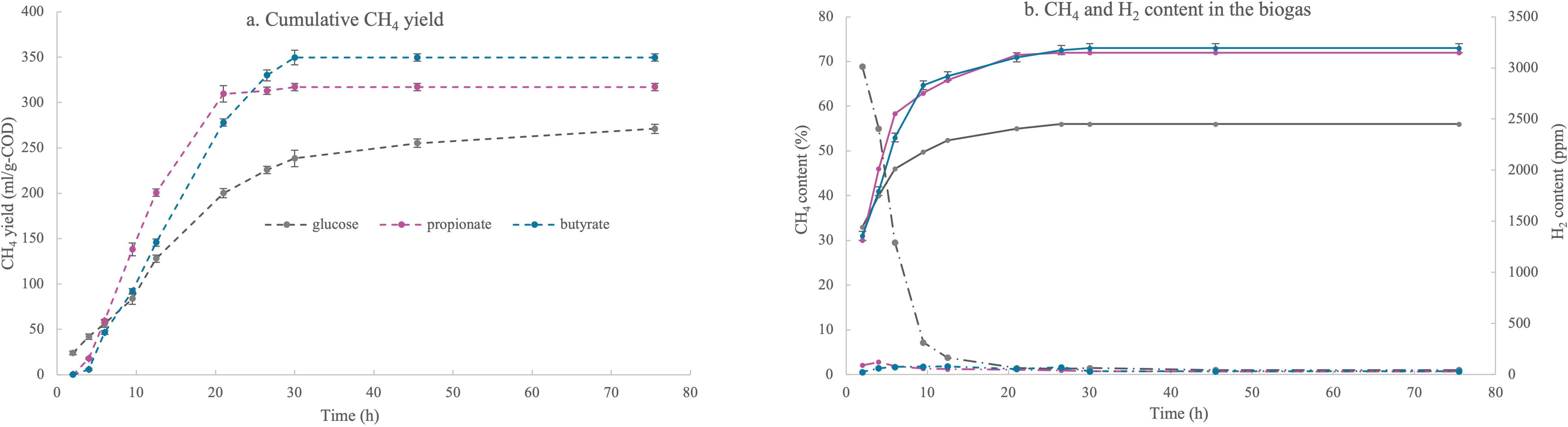
CH₄ yield and the CH₄ and H₂ contents in the biogas produced during the methanogenic conversion of glucose, butyrate, and propionate by the three enriched consortia on day 219.

**Figure 2.**
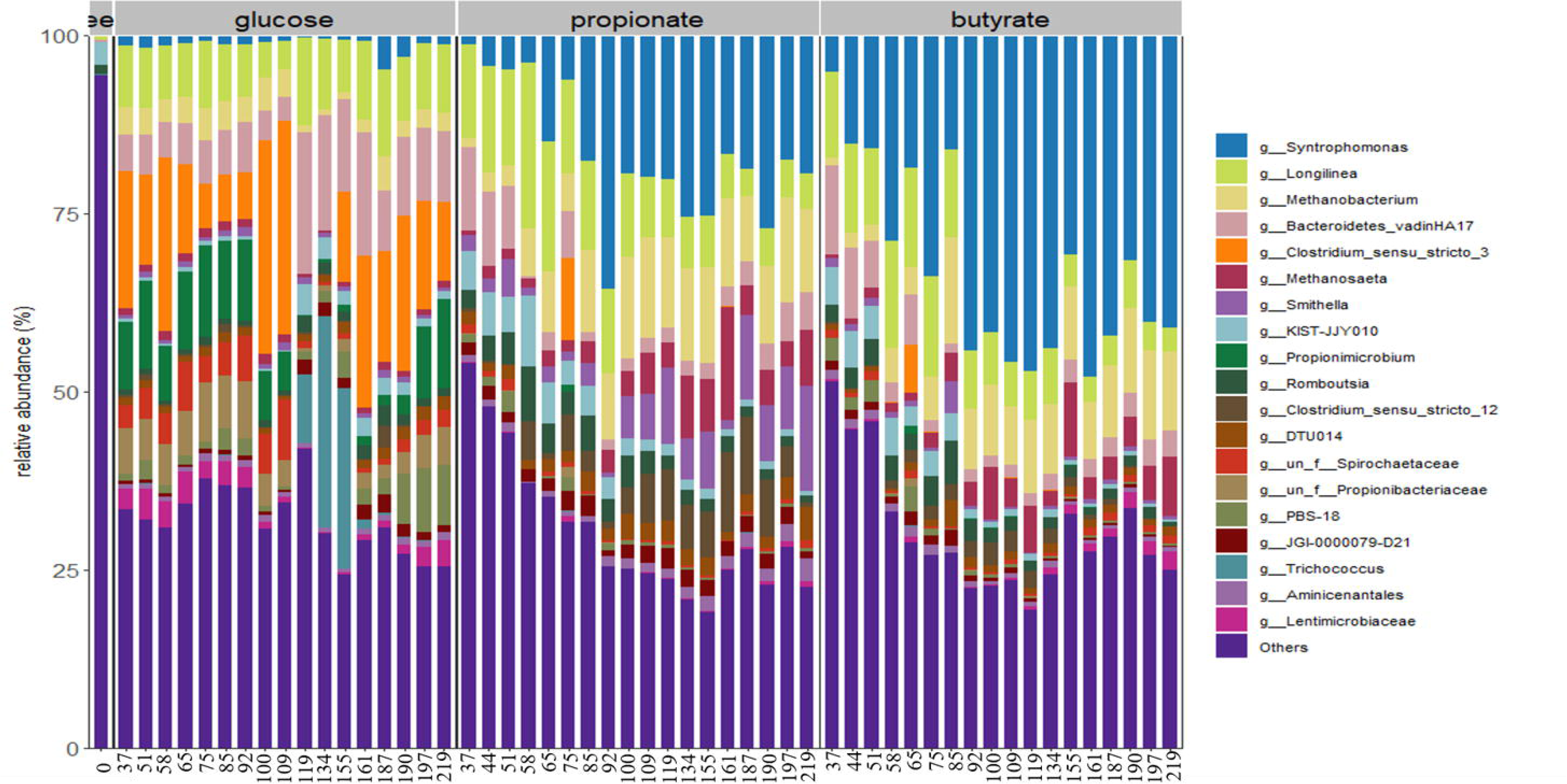
Taxonomy affiliation and relative abundance of the top 15 genus in the three consortia enriched with glucose, propionate and butyrate, respectively. Summarized in this figure are the 16S rRNA gene (v4-v5 region) based microbiome profile in Consortia-G, Consortia-B and Consortia-P from day 37 to day 219, and in seed sludge on day 0.

**Figure 3.**
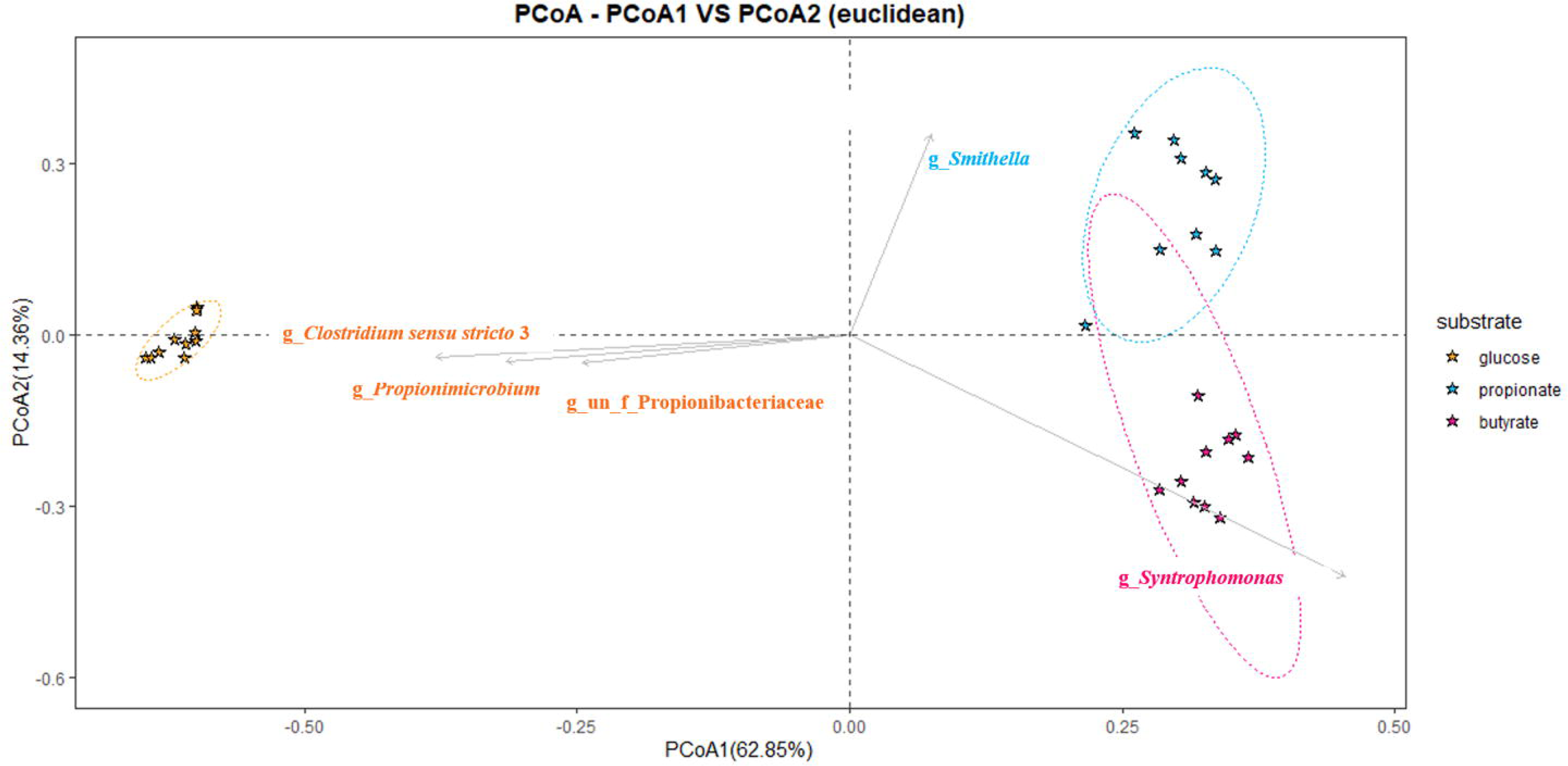
Principal coordinates analysis (PCoA) of the three anaerobic digestion consortia enriched with glucose, propionate and butyrate, respectively. The community structures were inferred from the 16S rRNA amplicon sequence variants (v4-v5 region), and the PCoA analysis is based on the euclidean distance matrix. PCoA1 and PCoA2 are shown with the percentage variation explained by each axis. During the course of the 219 days enrichment, the most varying population in Consortia-G were genus *Clostridium sensu stricto* 3, *Propionimicrobium*, and an unclassified genus in the family of *Propionibacteriacea*; the most varying population in Consortia-P was the genus *Smithella*, and the most varying population in Consortia-B was the genus *Syntrophomonas*.

We speculate that *Clostridium sensu stricto 3*, *Propionimicrobium*, and *f_Propionibacteriaceae* act as primary fermenting bacteria utilizing glucose, while *Smithella* functions as putative SPOB oxidizing propionate anaerobically, and *Syntrophomonas* as putative SBOB oxidizing butyrate anaerobically.

As summarized in Figure 2 and Figure S7-a, after 219 days of enrichment, the top 15 genera, which account for 75–83% of the populations in the three enriched consortia, exhibited only ∼4% relative abundance in the seed inoculum. Among these top 15 genera are two archaeal genera: *Methanosaeta* and *Methanobacterium*. Notably, the relative abundance of these methanogens, particularly *Methanobacterium* (Figure S7-b), is significantly higher in Consortia-B (20%) and Consortia-P (22%) compared to Consortia-G (6%).

### 3.2 Genome-centric transcriptome response of the primary fermenting bacteria, SBOBs and SPOBs to varying substrates

#### 3.2.1 Identification of MAGs associated with primary fermentation and syntrophic oxidation

In total, 317 bacterial MAGs are recovered from the 10 metagenome datasets. Summary of the taxonomy affiliation, relative abundance and transcription activity of each bacterial MAG can be found in Figure 4 and Appendix file 1. In the following genome-centric meta-omics analysis, we specifically focus on those 17 bacterial MAGs that have transcription response (>=0.5% of total mRNA clean reads mapped) to at least one of the substrates provided, i.e., glucose, propionate, butyrate, fumarate and crotonate. These 17 MAGs include FI21 and FI22 (hereafter, these two MAGs will be referred to as FI21, 22) affiliated to the genus of *Clostridium_T,* with the highest transcription activity towards glucose ; DES18, DES19, DES20, DES21, DES22, DES23, DES24, and DES25 (hereafter, these eight MAGs in the same genus of g_Fen_1166 will be referred to as DES18-25) affiliated to the family of *Smithellaceae*, with the highest transcription activity towards propionate; FI3, FI4 and FI7 (hereafter, these three MAGs will be referred to as FI3,4,7) affiliated to the genus of *Syntrophomoas_B,* with the highest transcription activity towards butyrate; ER1, ER2 and ER4 (hereafter, these three MAGs will be referred to as ER1,2,4) that are annotatable only at the phylum level as *Eremiobacterota*; and TM1 affiliated to the species *Mesotoga*. Considering that ER1, ER2, ER4, and TM1 exhibit inconsistent transcriptomic responses to the tested substrates, we hypothesize that they may function as fermentative opportunists within the consortia, survive on cell debris or other extracellular substances in the consortia. Further details on the functional niches of ER1,2,4 and TM1 are provided in the Supporting Information.

**Figure 4.**
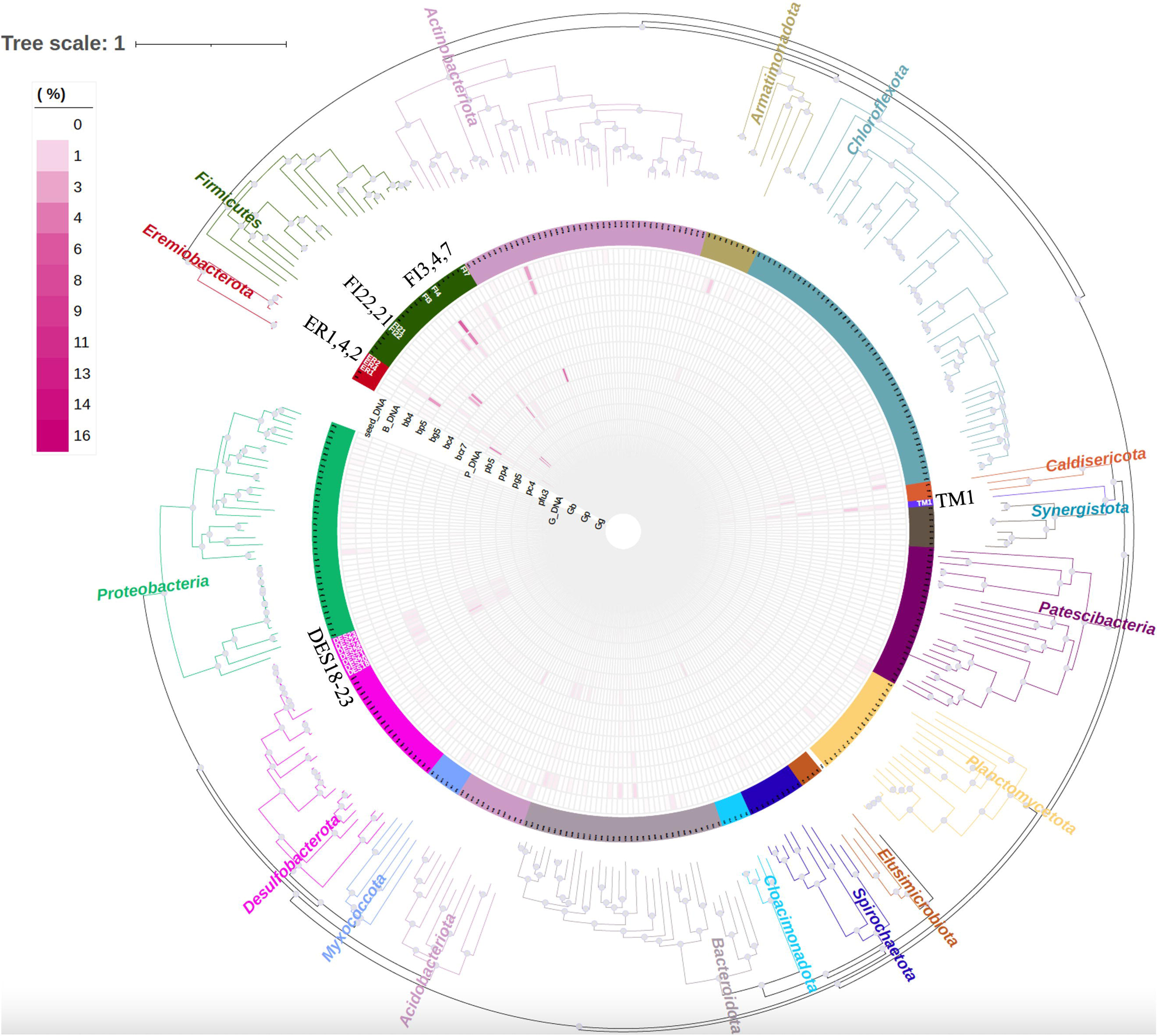
Maximum likelihood tree of the 315 bacterial MAGs recovered in this study. The inner heatmap summarized the meta-DNA based abundance and the meta-RNA based transcription activity of each MAG. ‘seed_DNA’, ‘B_DNA’, ‘P_DNA’ and ‘G_DNA’ denoted meta-DNA based abundance of each MAG in the seed sludge, in the enriched Consortia-G, Consortia-B and Consortia-P, respectively. The other datapoints are the meta-RNA based transcription activity of each MAG. Detailed information on each datapoint for meta-RNA based transcription analysis could be found in Table S1 and Figure S3- S5. The 17 MAGs highlighted with white bold labels were those MAGs annotated with relatively high transcription activity in response to certain substrates provided. This phylogenetic tree was constructed using FastTree [69] (maximum-likelihood phylogeny) based on a set of 120 concatenated universal single-copy proteins for Bacteria [41]. Branches of the phylogenetic tree are coloured according to their taxonomy affiliation at the phylum level, and those grey dots at each inner node indicate bootstrap values >=0.9.

The transcriptional activity of each functional gene in each MAG was analyzed and calculated independently, with a detailed summary of the RNA-RPKM values provided in Appendix file 1. In interpreting the transcription activity of bacterial MAGs in each of the three function guilds, as visualized in Figure 5, FI21, 22 were clustered to represent primary fermenting bacteria, FI3,4,7 was clustered to represent SBOBs, and DES18-25 were clustered to represent SPOBs. The MAGs within each of the three clusters are of ≥ 95% average nucleotide identity (or a pairwise mutation distance of ≤ 0.05, the cutoff used to group MAGs into same species by Mash [48]).

**Figure 5.**
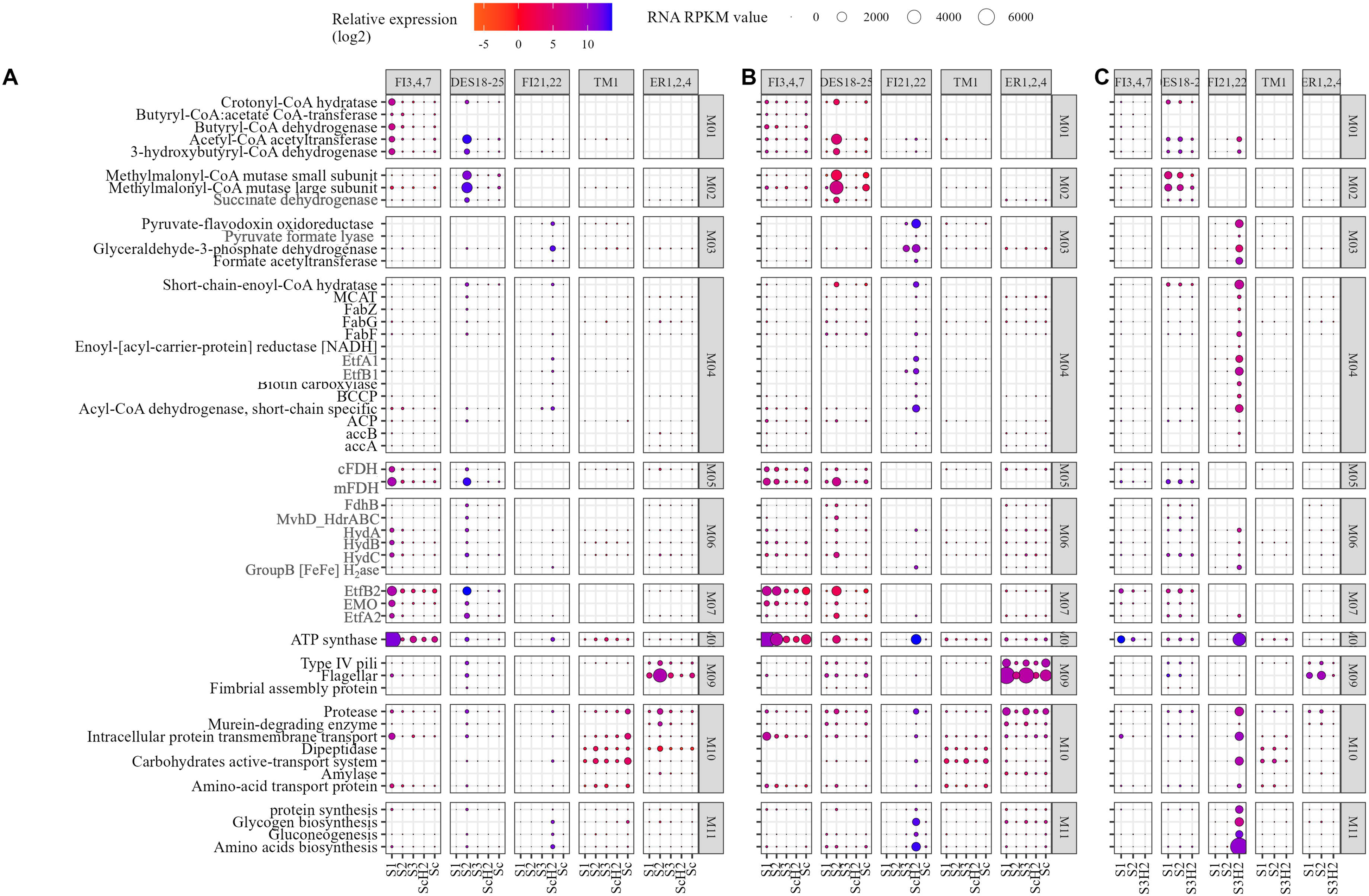
Variations in the transcription activity of the key functional genes annotated in the 17 MAGs in response to the stimuli of varying substrates in A) Consortia-B, B) Consortia-P and C) Consortia-G. FI3,4,7 refers to the three SBOB in the genera of *Syntrophomonas_B*, DES18-25 refers to the eight SPOB of a same unnamed genera in the family of Smithellaceae, FI21,22 refers to the two parimary fermenting bacteria in the genera of *Clostridium_T*, TM1 refers to the one *Mesotoga* MAG, and ER1,2,4 refers to the 3 co-occurring *Eremiobacterota* MAGs. S1, S2 and S3 listed horizontally denotated substrate as n-butyrate, propionate and fermentative substrate, respectively. The fermentative substrate applied is glucose in Consortia-G, fumarate in Consortia-P and crotonate in Consortia-B. Sc denoted substrate mix as glucose: propionate: butyrate=1:1:1 (COD: COD: COD). ScH_2_ and S3H_2_ refer to the datapoints when there is accumulation of H_2_, when fermentative substrate is present, or substrate mix is applied. Colors of the circles indicate changes in the transcription activity of the corresponding genes, in log difference relative to that of datapoint g0, b0 and p0 where no substrate is available at the end of an SBR batch. The size of the circles indicates the RNA-RPKM values of the corresponding genes. Clustered in M01 are key genes involved in butyrate oxidation, in M02 are key genes involved in propionate oxidation, in M03 are key genes involved in the glycolysis and pyruvate metabolism, in M04 are key genes involved in the fatty acid synthesis, in M05 are the cytosolic and membrane-bound formate dehydrogenase, in M06 are hydrogenase genes, in M07 are genes encoding the electron transfer flavoprotein. For clarity, the *EtfA* and *EtfB* subunits in the *Bcd_EtfAB_H_2_ase* gene cluster are denoted as *EtfA1* and *EtfB1*, while *EtfA* and *EtfB* subunits in the *Bcd_EtfAB_EMO* gene cluster are denoted as *EtfA2* and *EtfB2*. In M08 is the ATP synthase gene, in M09 are genes encoding the type IV pili, flagellar and fimbria assembly protein, in M10 are genes encoding the protease, murein-degrading enzymes and amylase etc., in M11 are genes encoding the synthesis of protein, amino acids, and glycogen. Full names of those genes denoted with abbreviations can be found in the Appendix file 1.

#### 3.2.3 Metabolic specificity of the primary fermenting bacteria, SBOBs and SPOBs

Metabolic specificity of anaerobes in the three function niches was clearly demonstrated by the transcription response of MAGs to varying substrates as summarized in Figure 5, and the substrate degradation profile as summarized in Figure S3, S4, S5. Specifically, when glucose was provided as the sole substrate to Consortia-G (datapoints gg2 in Figure 4), or as co-substrate to Consortia-B (datapoints bc4 in Figure 4) and Consortia-P (datapoints pc4 in Figure 4), FI21, 22 demonstrated the highest transcription activity in all the three consortia enriched, irrespective of their very low abundance in both Consortia-B and Consortia-P. Consortia-G enriched with FI21,22 cannot degrade either butyrate or propionate as rapid as Consortia-B or Consortia-P, probably due to low abundance of the SBOB and SPOB populations in the community, and the enriched FI21,22 are of poor metabolic capacity towards either butyrate or propionate as indicated by their low transcription response to either butyrate or propionate (Figure 4). When butyrate was provided as substrate to Consortia-B (datapoint bb4 in Figure 4) or to Consortia-P (datapoint pb5 in Figure 4), FI3,4,7 are noted with the highest transcription activity in both consortia. When propionate was provided as the primary substrate to either Consortia-B (datapoint bp5 in Figure 4) or Consortia-P (datapoint pp4 in Figure 4), consistent and relatively high transcription activity of DES18-25 is observed in both consortia.

### 3.3 Transcription activity of function gene clusters mediating either flavin based electron bifurcation (FBEB) or reverse electron transfer (RET) in primary fermentation, in anaerobic butyrate oxidation and in anaerobic propionate oxidation

Summarized in Table 1 are the genes/gene clusters mediating either the flavin-based electron bifurcation (FBEB) or reverse electron transfer (RET) that are the featuring biochemistry in primary fermentation, or in syntrophic butyrate and propionate oxidation.

**Table 1.** Presence (+) and absence(-) of the gene clusters mediating either flavin based electron bifurcation (FBEB) or reverse electron transfer (RET) in primary fermentation and in syntrophic oxidation of butyrate and propionate, their cellular component and the corresponding biochemistry

| Gene/ Gene clusters | Cellular component | Biochemical reaction | Primary fermenting bacteria | SBOB | SPOB |
| --- | --- | --- | --- | --- | --- |
|  |  |  | FI21,22 | FI3,4,7 | DES18-25 |
| *hydA_hydB_hydC* | Cytoplasm | a.1) $\text{Fd}_{\text{red}} + \text{NADH} + 4\text{H}^+ = 2\text{H}_2 + \text{NAD}^+ + \text{Fd}_{\text{ox}}$<br>(in primary fermentation);<br>a.2) $\text{NADH} + 2\text{H}^+ = \text{H}_2 + \text{NAD}^+$ (in syntrophic oxidation) | + | + | + |
| *Bcd_EtfA_EtfB_H<sub>2</sub>ase* | Cytoplasm | b) $2\text{NADH} + \text{Crotonyl-CoA} + \text{Fd}_{\text{ox}} = 2\text{NAD}^+ + \text{Butyryl-CoA} + \text{Fd}_{\text{red}}$ | + | - | - |
| *cFDH* | Cytoplasm | c) $\text{NADH} + \text{CO}_2 + \text{H}^+ = \text{NAD}^+ + \text{HCOOH}$ | - | + | + |
| *mFDH* | Membrane-bound | d) $(\text{M})\text{MKH}_2 + \text{CO}_2 + \text{PMF} = \text{HCOOH} + (\text{M})\text{MK}$ | - | + | + |
| *FdhB_mvhD_HdrABC* | Cytoplasm | UNKNOWN | - | - | + |
| *Bcd_EtfB_EMO_EtfA* | Membrane-bound | e) $\text{Butyryl-CoA} + \text{MMK} + \text{PMF} = \text{Crotonyl-CoA} + \text{MMKH}_2$ | - | + | - |
| *SdhA_SdhB_SdhC* | Membrane-bound | $\text{Succinate} + (\text{M})\text{MK} + \text{PMF} = \text{Fumarate} + (\text{M})\text{MKH}_2$ | - | - | + |

#### 3.3.1 The FBEB type hydABC gene cluster in primary fermenting bacteria and the non- FBEB type hydABC gene cluster in syntrophic oxidizing bacteria

High transcription activity of the cytosolic FBEB *HydABC* gene cluster in FI21,22 suggests the involvement of electron confurcation of Fd_red_ and NADH for H_2_ generation during acetate generation (Figure 6a) in primary fermentation of the AD bioprocess where hydrogenotrophic methanogens keep the H_2_ concentration low.

**Figure 6.**
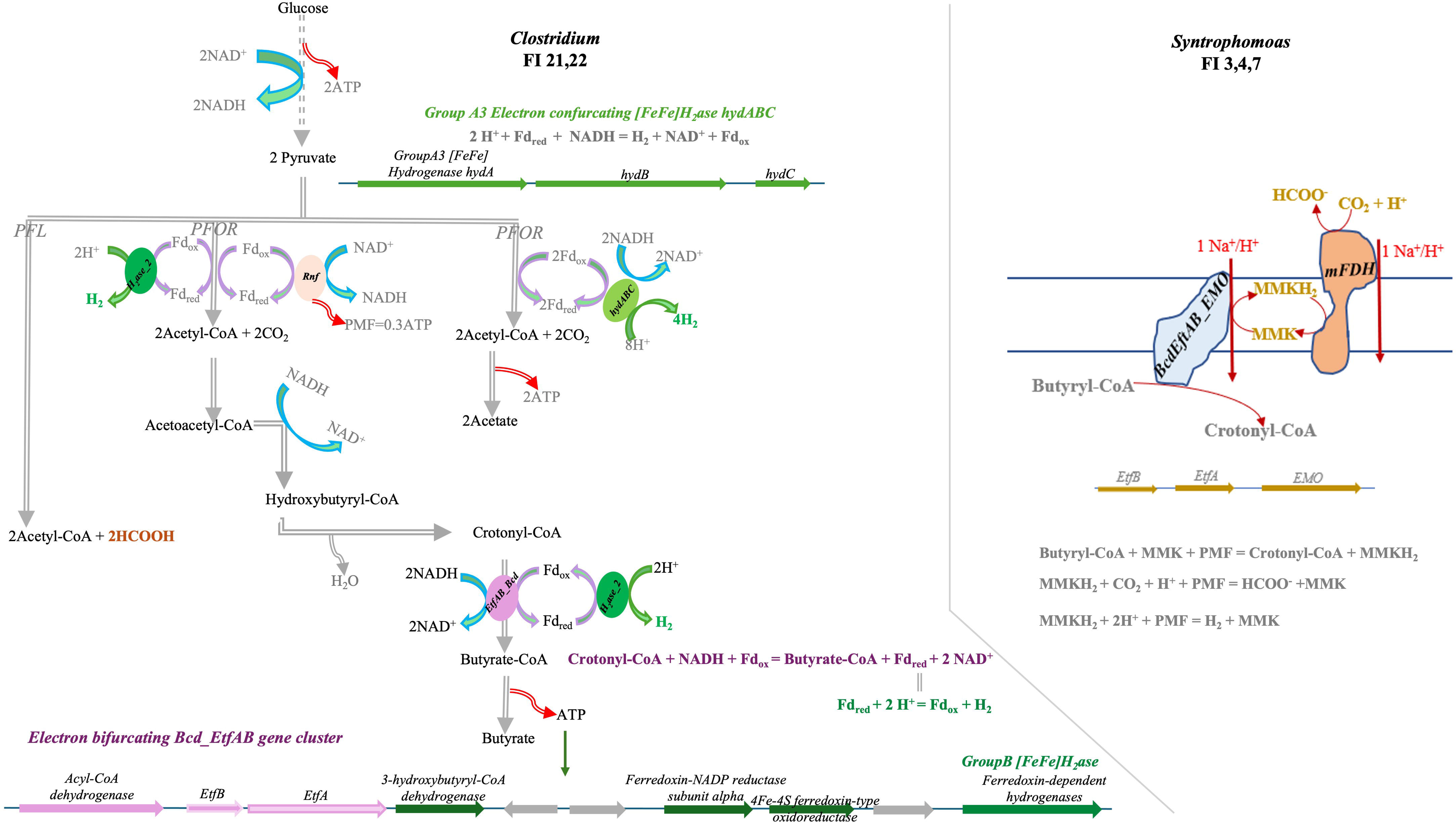
a) Metabolic pathways involved in H2 and formate generation during primary fermentation; c) RET-meditated formate and/or H2 generation during anaerobic oxidation of butyrate.

The *hydABC* gene cluster is also annotated and with transcription activity in both SBOB *Syntrophomoas* FI3,4,7 and in SPOB *Smithelacea* DES18-25 (Figure 5). However, as is visualized in Figure 7, comparing with that in the primary fermenting bacteria, mutation at the C-terminal end of *HydB*, which is a lack of the 2*[4Fe-4S] function domains, is noted in both *Syntrophomoas* FI3,4,7 and in *Smithelacea* DES18-25. Same mutation has also been reported in the *hydB* subunit of *S. aciditrophicus* [49] and of *S. wolfei* [50]. Biochemical characterization revealed that enzyme products of the *hydABC* gene cluster in both *S. aciditrophicus* and *S. wolfei* cannot interact with Fdred and can use NADH as direct electron donor for the reduction of H^+^ [49] [50]. NADH-coupled H⁺ reduction in syntrophs is thermodynamically feasible only under sufficiently low H₂ concentrations (2.2–18.0 Pa) [49, 50] [51]. In other words, when H₂ concentrations are high, H⁺ as an electron acceptor for NADH reoxidation becomes unfavorable. Under such conditions, CO₂ may serve as an alternative electron acceptor for NADH reoxidation, leading to the production of formate, provided the formate concentration remains low.

**Figure 7.**
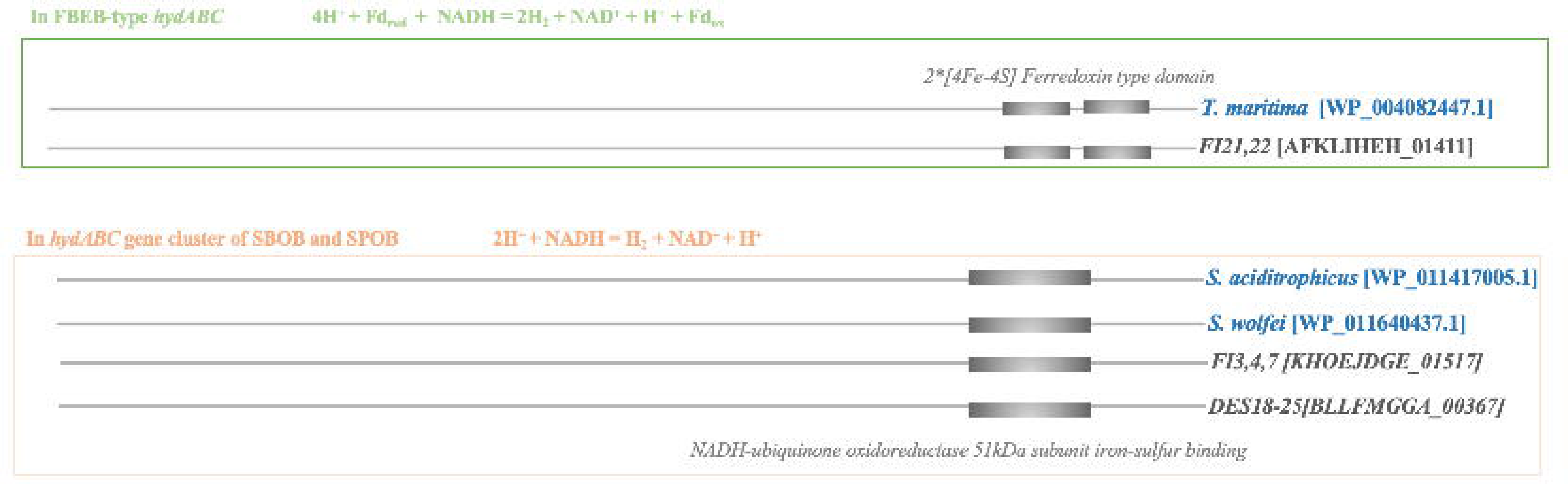
Summary of the C-terminal end domain in the *hydB* subunit in the flavin <u>b</u>ased <u>e</u>lectron <u>b</u>ifrucation (FBEB) type of *hydABC* in the primary fermenting populations (upper box) and in the non- FBEB type *hydABC* in the SBOB and SPOB. Name of the reference microbe strains in which the catalytic activity of the corresponding *hydABC* enzyme complex has been characterized are highlighted in blue. The names of the anaerobe populations enriched in study are shown in grey. Provided in the brackets are the gene accession number of the corresponding *hydB* gene.

#### 3.3.2 Cytosolic and membrane-bound formate dehydrogenase in syntrophic oxidizing bacteria

Gene clusters encoding formate dehydrogenase (FDH) are uniquely annotated and identified with transcription activity in the secondary fermenting populations enriched in this study, i.e., SBOB *Syntrophomoas* FI3,4,7 and SPOB *Smithelacea* DES18-22.

The cytosolic formate dehydrogenase (*cFDH*) catalyzes NADH-coupled HCO_3-_/CO_2_ reduction and generates low concentrations of intracellular HCOOH/HCOO^-^. The membrane-bound formate dehydrogenase (*mFDH*) catalyzes the proton motive force (PMF)-dependent reverse electron transfer (RET) during the endergonic CO_2_ reduction with the membranous (methyl)menaquinone (M)MKH_2_, and generates low concentration of formate extracellularly. During syntrophic propionate and butyrate oxidation, low concentrations of formate maintained by both efficient interspecies formate transfer [52, 53] and by activity of the partnering formate-utilizing methanogen is essential for such formate-centered recycling of the redox cofactors: NAD^+^ and (M)MK^+^ [54].

In comparison, neither *cFDH* nor *mFDH* is annotated in genomes of the primary fermenting bacteria *Clostridium* FI21,22, therefore, formate is not relevant in the recycling of redox cofactors (Fd_ox_ or NAD^+^) during primary fermentation. Formate as a product of primary fermentation can originate from the activity of the pyruvate formate lyase (*PFL*), which biochemistry is not associated with the recycling of the redox cofactors.

Formate hydrogenlyase (*FHL*) is not annotated in either the primary fermenting populations or the secondary fermenting populations. Under physiological conditions, bacterial *FHL* can mediate the unidirectional conversion of formate to H_2_, and its absence suggests the lack of biochemistry for the direct interconversion of formate and H_2_ in the key functioning bacteria enriched in this study. In other words, metabolism of the two dominant electron carriers, H_2_ and formate, are segregated in both primary fermenting bacteria, in SBOBs and SPOBs enriched in this study.

#### 3.3.3 The cytosolic Bcd_EtfAB_H_2_ase gene cluster in primary fermenting bacteria and the membrane-bound Bcd_EtfAB_EMO gene cluster in syntrophic butyrate oxidizing bacteria

Butyryl-CoA dehydrogenase (*Bcd*) genes clustering with the electron transfer subunits *EtfA* and *EtfB* (hereafter referred to as *Bcd_EtfAB*) exhibit high transcriptional activity in both the primary fermenting bacterium Clostridium FI21,22 and the SBOB Syntrophomonas FI3,4,7. The *Bcd_EtfAB* gene clusters in primary fermenting bacteria and in SBOBs can be distinguished by their associated neighboring genes, being either a Group B ferredoxin-dependent [FeFe] hydrogenase (H₂ase) or a membrane-bound oxidoreductase (*EMO*).

In Clostridium FI21,22, the Bcd_EtfAB cluster neighbors a Group B H₂ase (Figure 6a). This *Bcd_EtfA1B1_H₂ase* gene cluster mediates two key reactions: 1) NADH-dependent FBEB biochemistry for the reduction of crotonyl-CoA to butyryl-CoA (Reaction ‘b’ in Table 1) [55], and 2) Subsequent reduction of protons (H⁺) to hydrogen gas (H₂), driven by ferredoxin reduction (Fd_red_) (Reaction ‘c’ in Table 1) [56].

In contrast, the *Bcd_EtfAB* in genomes of SBOB populations FI3,4,7 cluster with a *EMO* subunit. Unlike the cytosolic *Bcd_EtfAB_H₂ase* enzyme complex, this *Bcd_EtfAB_EMO* enzyme complex is membrane-associated and catalyzes the proton motive force (PMF)- dependent butyryl-CoA oxidation mediated by membrane-bound methylmenaquinone (MMK) (Figure 6b, and Reaction ‘b’ in Table 1) [57]. The MMKH₂ produced in this step is reoxidized in the engergonic CO₂ reduction to formate, catalyzed by either a membrane-bound formate dehydrogenase (*mFDH*) or *mH_2_ase* [57].

#### 3.3.4 Anaerobic succinate oxidation and the FdhB_mvhD_hdrABC gene cluster in syntrophic propionate oxidizing Smithellaceae

The SPOB populations, DES18-25, that were specifically enriched in this study belong to an undefined genus within the Smithellaceae family. Propionate conversion via the dismutation pathway has been reported only in *Smithella propionica*, where two propionate molecules are first converted to acetate and butyrate, and the butyrate is subsequently converted via β- oxidation [58, 59]. In contrast, the methylmalonyl-CoA pathway for propionate degradation has been identified in all other SPOBs characterized to date [5].

The SPOB DES18-25, enriched in this study, is part of the Smithellaceae family, though the resolution of the taxonomy is not sufficient to confirm whether it is of the genus *Smithella propionica*. As summarized in Figure 5, both the presence and transcriptional activity of genes involved in β-oxidation of butyrate and in the methylmalonyl-CoA pathway have been observed in Smithellaceae DES18-25.

One of the interesting transcription patterns noted in DES18-25 is of the *FdhB_MvhD_HdrABC* gene cluster. Transcription activity and proteome signal of the *FdhB_MvhD_HdrABC* gene cluster in SPOBs have been reported in previous studies [60, 61]. Biochemical characterization of the *FdhAB_mvhD_HdrABC* enzyme complex in methanogens revealed their FBEB activity during methanogenesis from formate [54], while explicit biochemical evidence is absent regarding the catalytic activity of the *FdhB_MvhD_HdrABC* enzyme complex commonly annotated in SPOBs. Currently, there are two hypotheses regarding the function roles of this bacterial *FdhB_MvhD_HdrABC* enzyme complex: it may either be involved in sulfate reduction [61] or may possess an uncharacterized FBEB activity [60]. We tentatively favor the second hypothesis for two reasons: 1) it was reported that the gram-negative bacteria *Smithella* cannot reduce sulfate [5], further, in this study, to exclude the interference of sulfate as electron acceptor, MgSO_4_·7H₂O was replaced with MgCl₂·6H₂O (Supporting information); and 2) the *FdhB_MvhD_HdrABC* gene cluster showed high transcriptional activity almost exclusively under the presence of propionate as substrate.

### 3.4 Segregation of H_2_ and formate metabolism in primary and secondary fermentation

As is discussed in the above section, in primary fermenting bacteria FI21,22, FBEB type *hydABC* and ferredoxin-dependent [FeFe] H_2_ase are annotated and transcribed, mediating electron transfer to H_2_ during primary fermentation, and no formate dehydrogenase (*FDH*) was annotated or transcribed. In contrast, uniquely transcribed with high activity in the secondary fermenting populations, i.e., SBOBs FI3,4,7 and SPOBs DES18-22, are *FDH*. Interconversion of H_2_ and formate is limited in these bacteria due to the absence of the *FHL*.

One of the ecological relevance of such segregation of the H_2_ and formate metabolism could be that the ‘H_2_-tolerant’ primary fermentation and the ‘H_2_-sensitive’ secondary fermentation can proceed simultaneously, with H_2_ being the dominant electron sink in the former, and formate as the predominant electron sink in the latter. Our wet-lab observation aligns well with this hypothesis. Accumulation of H_2_ in high concentrations (5,000-70,000 ppm, Figure S8) was observed only when fermentative substances (i.e., glucose and fumarate) are present as either sole substrate or as co-substrate. H_2_ accumulation was never detected when only either propionate or butyrate was supplied as the primary substrate. H₂ concentrations in a much lower range of 4.6-18.0 Pa (46–180 ppm) [50] and 6.8 Pa (∼70 ppm) [62] during syntrophic metabolism has also been reported before.

If we assume that H₂ is the sole electron carrier during secondary fermentation, the acetogenic oxidation of propionate and butyrate would become thermodynamically unfavourable at gaseous H₂ concentrations exceeding 132 ppm and 134 ppm, respectively (Figure S9)^1^. However, contrary to this theoretical calculation, when a substrate mixes of glucose: propionate: butyrate (COD ratio 1:1:1) was provided, intermittent H₂ accumulation exceeding 30,000 ppm was detected in both Consortia-B and Consortia-P—far higher than the theoretical inhibition thresholds for H₂. Despite such high H₂ concentrations, propionate and butyrate degradation were not completely inhibited, as evidenced by their declining concentrations over time (Figure 8).

**Figure 8.**
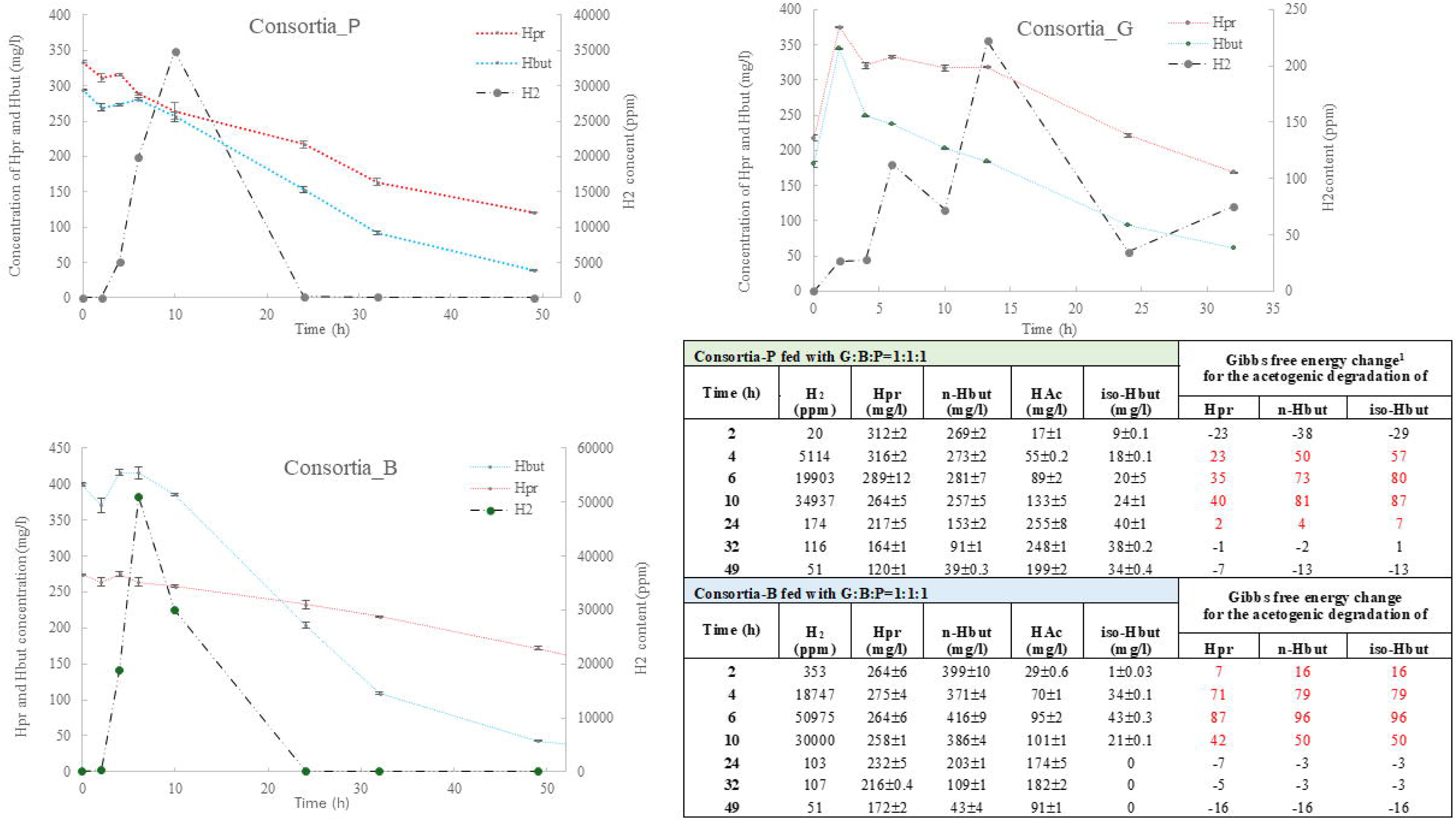
Degradation profile of propionate (HPr) and butyrate (HBut), and dynamics of H_2_ concentration when substrate mix as glucose:propionate:butyrate = 1:1:1 (COD:COD:COD) was provided as substrate in a) Consortia-P, b) Consortia-B, and c) Consortia-G. d) Gibbs free energy change calculated for the degradation of propionate, n-butyrate and iso-butyrate in Consortia-P and Consortia-B , respectively, assuming that H_2_ is the only electron carrier generated from the syntrophic oxidation of butyrate or propionate.

Notably, studies from the 1980s reported H₂ inhibition concentation of 2.03 kPa for anaerobic butyrate oxidation and H_2_ inhibition constant at 11.1 kPa for anaerobic propionate oxidation [29, 30]. However, these findings have been largely overlooked, and no follow-up research has addressed why propionate and butyrate oxidation can proceed under such high H₂ concentrations. Combining with the observations in this study, one plausible explanation could be that formate, rather than H₂, acts as the dominant interspecies electron carrier during syntrophic oxidation. Butyrate and propionate oxidation can remain active under elevated H₂ concentrations if there is no available biochemistry to mediate a direct conversion of H_2_ to formate. Formate concentration at low levels is essential for such formate-driven syntrophic metabolism, which can be maintained with efficient interspecies formate transfer and rapid formate utilization by the coupling methanogens in the enriched consortia [54].

### 3.5 Formate- and H_2_-utilizing methanogen and acetoclastic methanogen identified in the enriched consortia

Among the 39 archaeal MAGs recovered (Figure 9), 4 archaeal MAGs are of the highest transcriptional activity, accounting for (79±9) % of the mRNA gene transcripts mapped to the 39 archaeal MAGs. These 4 archaeal MAGs are: EA7 affiliated with the family of *Methanobacteriaceae*, and HA1, HA2, HA3 (hereafter, termed as HA1,2,3) affiliated with the species of *Methanothrix soehngenii*. Therefore, in the genome-centric interrogation, we focused specifically on these 4 archaeal MAGs to elaborate on the methanogenic activity of the enriched consortia.

**Figure 9.**
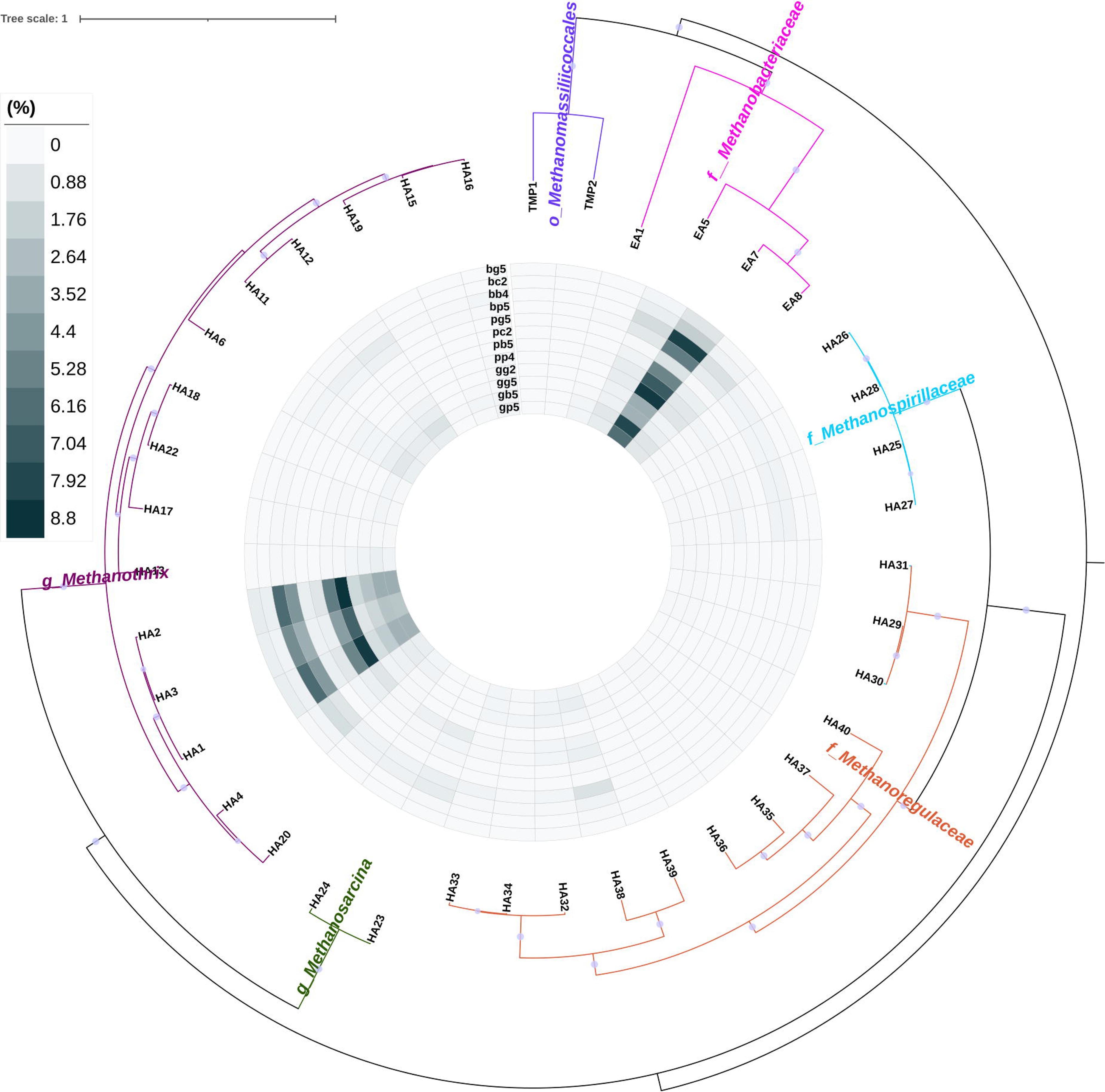
Maximum likelihood tree of the 39 archaeal MAGs recovered in this study. The inner heatmap summarized the meta-RNA based transcription activity of each MAG. Detailed information on each meta-RNA datapoint could be found in Table S1. The tree was constructed using FastTree (maximum- likelihood phylogeny) [69] based on a set of 122 concatenated universal single-copy proteins for *Archaea* [41]. Branches of the phylogenetic tree are coloured according to their taxonomy affiliation, and those light purple dots indicate bootstrap values >=0.9.

According to the transcription profile as summarized in Figure 10 and Figure S10, EA7 is a formate- and H_2_- utilizing methanogen and HA1,2,3 are acetoclastic methanogens. Specifically, *Methanobacteriaceae* EA7 dominated the transcription activity of genes encoding Formate dehydrogenase (coenzyme F420-coupled, denotated as *FdhAB*), formylmethanofuran dehydrogenase (*fwd*), and coenzyme F420 hydrogenase (*frh*). In comparison, *Methanothrix soehngenii* HA1,2,3 dominated the mRNA transcripts of genes encoding the acetyl-CoA decarbonylase (*cdhDE*) and the carbon-monoxide dehydrogenase (*cdhAB*). The acetoclastic archaeal MAGs HA1,2,3 are of high identity to strain *Methanothrix soehngenii*, while the hydrogenotrophic archaeal MAG EA7 are annotatable only at the family level of *Methanobacteriacea,* indicating novelty of this hydrogenotrophic methanogen population enriched.

**Figure 10.**
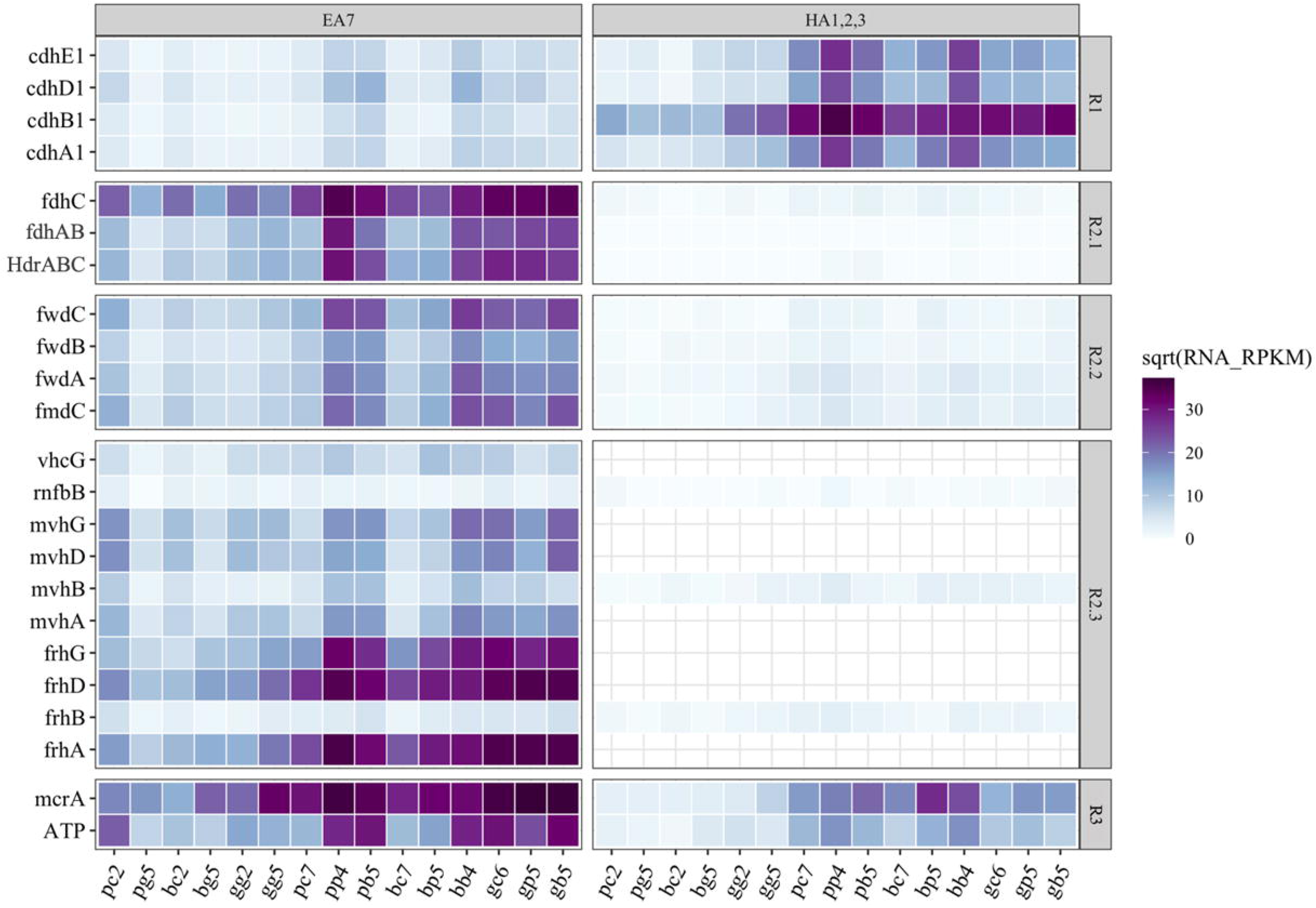
Transcription activity of the key functional genes annotated in the Methanobacteriaceae MAG EA7, and in the three *Methanothrix* MAGs HA1,2,3. The heatmap represents the transcription activity of the corresponding genes, and the data given alongside the heatmap bar are square root of the RNA-RPKM values. Full names of genes denoted with abbreviations can be found in the Appendix file 1. Genes grouped in grid R1 are those specifically involved in acetoclastic methanogenesis, genes grouped in grid R2.1-R2.3 are those specifically involved in hydrogenotrophic methanogenesis, and genes grouped in grid R3 are genes shared by both the hydrogenotrophic methanogenesis and acetoclastic methanogenesis

Formate as an important electron carrier in the syntrophic cooperation between the SBOB/SPOB and the methanogen archaea is hinted also by the high transcription activity of the *fdhC* gene encoding formate transporter in EA7, and also by the presence and the high transcription activity of the *FdhAB_mvhD_HdrABC* gene cluster. The formate-centered FEBE biochemistry mediated by this *FdhAB_mvhD_hdrABC* enzyme complex is reaction (1). Also annotated in EA7 is the presence and transcription activity of the *MvhAG_MvhD_HdrABC* gene cluster, mediating the H_2_-centered FBEB reaction (2). Both *MvhAG_MvhD_HdrABC* and *FdhAB_mvhD_hdrABC* can couple the first step and last step of wolf cycle in methanogenesis[63].

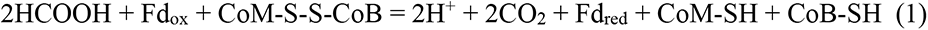

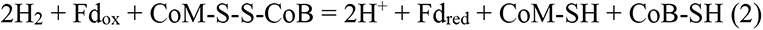

### 3.6 Metabolic specificity of the key functional populations in the AD consortia

This study reveals a clear functional partitioning of the various metabolic processes in anaerobic digestion, including primary fermentation, syntrophic butyrate oxidation, syntrophic propionate oxidation, hydrogenotrophic methanogenesis, and acetoclastic methanogenesis, each carried out by distinct microbial populations. As illustrated in Figure 11, the characteristic gene clusters underlying these biochemistries are assigned to the corresponding anaerobic populations.

**Figure 11.**
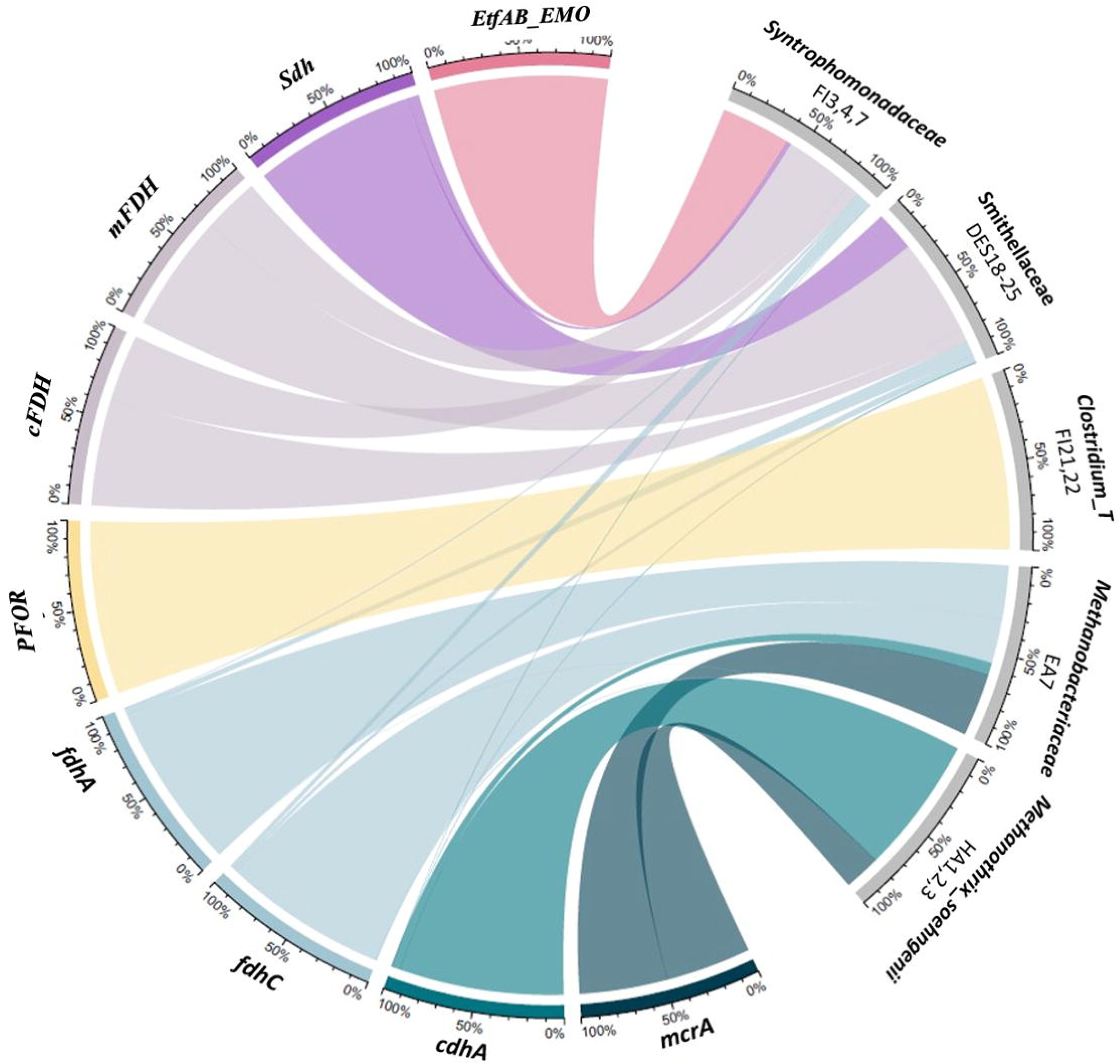
Contribution of the putative fermenter bacteria *Clostridium_T* FI21,22, the SBOB *Syntrophomonadaceae* FI3,4,7), the SPOB *Smithellaceae*, DES18-25 to the mRNA transcripts of the core genes (left, coloured by genes) involved in the butyrate metabolism (*EMO, membrane-bound oxidoreductase clustering with Bcd_EtfAB*), the propionate metabolism (*Sdh*, Succinate dehydrogenase) and the pyruvate fermentation (*PFOR,* Pyruvate-ferredoxin oxidoreductase), in response to the availability of butyrate, propionate and glucose, respectively. And also, contribution of *Methanobacteriaceae* EA7 and *Methanothrix_soehngenii* HA1,2,3 to the mRNA transcripts of the core genes involved in the methanogenesis metabolism, i.e., *fdhA*: formate dehydrogenase (coenzyme F420) alpha subunit, *fdhC*: formate transporter, *cdhA*: acetyl-CoA decarbonylase, and *mcrA*: methyl coenzyme M reductase. Colours represent different genes, and the width of each ribbon represents the contribution ratios of each microbe group to the mRNA transcripts of each gene, the ratios were averaged across the transcription activity of the corresponding populations in the three enriched consortia: Consortia-G, Consortia-B and Consortia-P. *cFDH* and *mFDH* refer to the cytosolic and membrane-bound formate dehydrogenase, respectively.

In all three consortia enriched, when butyrate was the primary or co-substrate, 100% of the *EMO* gene transcripts in the *Bcd_etfAB_EMO* gene cluster were expressed by the SBOBs. For propionate as the primary or co-substrate, 90±5% of the *Sdh* gene transcripts were expressed by the SPOBs. When glucose was provided, the primary fermenting populations contributed 97±2% of the transcripts for genes involved in glycolysis and fermentation metabolism, such as the *PFOR* gene.

Formate dehydrogenase genes, including *cFDH* and *mFDH*, were exclusively transcribed in syntrophic populations, i.e., SBOBs (FI3,4,7) and SPOBs (DES18-22). Coenzyme F420- dependent formate dehydrogenase (*FdhAB*) was exclusively transcribed in the hydrogenotrophic methanogen EA7, highlighting the role of formate as the electron carrier between syntrophic oxidizers and methanogens.

The two groups of methanogens, Methanothrix (HA1,2,3) and Methanobacteriaceae (EA7), equally contributed to overall CH₄ generation. Both groups each contributed ∼50% of the transcripts for Methyl-coenzyme M reductase (*mcrA*). Specifically, Methanobacteriaceae EA7 dominated the conversion of formate and H₂, contributing 95±2% of the relevant transcripts (e.g., *frhA, fdhC* genes). Meanwhile, Methanothrix soehngenii HA1,2,3 dominated the conversion of acetate, contributing 92±4% of the transcripts for functional genes, such as *cdhA*. Additionally, we assessed the metabolic flexibility of SBOBs and SPOBs by testing their ability to utilize crotonate or fumarate. At the community level, crotonate fermentation for butyrate generation was detected in Consortia-B (Figure S3) and fumarate fermentation for propionate generation in Consortia-P (Figure S4), following a very long lag phase of 40–50 hours. However, due to the low transcription activity of FI3,4,7 and DES18-25 for these fermentative substrates (denoted as S3 in Figure 5), we could not conclude that SBOBs or SPOBs are responsible for crotonate or fumarate fermentation via the reverse syntrophic oxidation of butyryl-CoA or succinate. This suggests that the Syntrophomonas_B SBOB (FI3,4,7) and Smithellaceae SPOB (DES18-25) populations enriched in this study exhibit limited metabolic flexibility towards utilizing fermentative substrates.

### 3.7 Limitations of this study

In this study, through enrichment experiments, batch tests, and the integration of physicochemical parameter measurements with intensive genome-centric transcriptome profiling, we identified and characterized populations within key functional guilds of the AD consortia. The key findings have been summarized and elaborated above. Nevertheless, we want to address several important limitations of this study, which will guide future research, either by us or by other researchers working in this field. These limitations include: 1) the lack of direct formate measurements; 2) the unaddressed potential for chain elongation and direct electron transfer (DIET), which were not directly evaluated; and 3) the absence of mass balance analysis.

#### 3.7.1 Lack of direct measurement of formate

Formate was not directly measured in this study and formate as the electron carrier mediating syntrophic oxidation under higher H₂ concentrations is indirectly inferred from thermodynamic evaluations and from the transcriptional activity of formate transporters and formate dehydrogenase in SPOBs, SBOBs, and methanogens.

Literature on formate as a major interspecies electron carrier in syntrophic metabolism often are based also on indirect evidence—such as elevated levels of *FDH* [32], or formate accumulation induced by methanogenesis inhibition (e.g., CHCl₃ treatment) [64], or halted syntrophic oxidation in mutant methanogens deficient in formate utilization [65].It is rare that formate as dominant electron carrier in syntrophic metabolism is inferred from a direct measurement, which largely is due to the fact that, with its high solubility and diffusion kinetics, accumulation or detectable concentration of formate is not expected in such metabolism [66]. Nevertheless, it would strengthen our study to get formate measured directly, even if it may fall below detectable levels. We attempted in this study to measure formate using GC-FID but encountered challenges, as the FID response for formic acid is very low and therefore not useful, especially for the measurement of lower concentrations of formic acid. In future studies, we will work with more sensitive HPLC methods to yield more definitive results.

#### 3.7.2 Lack of direct evaluation on the relevance of chain elongation or DIET in butyrate and propionate degradation under high H_2_ concentrations

Previous studies have highlighted the role of chain elongation in converting butyrate or propionate to caproate or valerate under high H₂ concentrations [67]. However, direct measurements of caproate and valerate were not conducted in this study, and we tentatively exclude chain elongation based on the following indirect evidence: (1) among the key enzymes for ethanol fermentation (ethanol is an essential precursor for chain elongation), only alcohol dehydrogenase was annotated in the primary fermentative bacteria FI21 and FI22, and its transcription activity was low (Appendix File 1); and (2) no transcriptional activity of genes essential for butyrate or propionate chain elongation was observed. Specifically, genes involved in butyrate elongation (e.g., hexanoyl-CoA synthase, hydroxyhexanoyl-CoA oxidoreductase) and propionate elongation (e.g., pentanoyl-CoA synthase, hydroxypentanoyl-CoA oxidoreductase) were absent.

Similarly, while DIET has been reported to enable anaerobic VFA oxidation at high H₂ partial pressures [68], we did not evaluate its relevance directly. Indirect evidence suggests limited involvement: (1) DIET typically involves *Geobacter* species, which were not annotated in our consortia; (2) no conductive carbon materials were used in this study, and the sludge appeared as loose flocs, not dense granules; and (3) no transcriptional activity was observed for genes associated with DIET, specifically *pilA*, *pilR*, *omcS*, *omcT*, or *recA*. A detailed summary of transcription activity for non-e-pili type IV pili genes in SBOBs and SPOBs is included in Appendix File 1, but none correspond to known DIET-relevant pathways.

#### 3.7.2 Insufficient quantitative insights

Although we aimed to achieve robust quantitative insights in this study, we eventually gave up in establishing the desired mass balance analysis, due to the limited reactor and equipment resources available; and the results we obtained are primarily descriptive. Ideally, each sampling point in the batch tests would include a comprehensive mass balance analysis, accounting for substrates and a full spectrum of metabolic products in both the gaseous and liquid phases. Additionally, determining specific H₂/formate inhibition concentrations or inhibition constants would have added quantitative rigor. However, such analyses were not feasible with our equipment set. We hope future efforts, either by us or other researchers, will provide these critical quantitative insights.

## Data Availability Statement

Metagenome DNA (10 datasets in total) and metatranscriptome RNA (20 data sets in total) sequencing data are available in the NCBI Sequence Read Achieve (SRA) database under the Bioproject accession number of PRJNA886957. The 17 bacterial MAGs and 4 archaeal MAGs analyzed in this study are accessible in NCBI under the Bioproject accession number of PRJNA886046, function genes annotated in these 21 MAGs and their transcription activity could be found in Appendix file 1. 16S rRNA gene amplicon sequences (v4-v5 region) have also been deposited in NCBI under the Bioproject accession number of PRJNA885972.

## Abbreviations

SBOBs - syntrophic butyrate oxidizing bacteria; SPOBs - syntrophic propionate oxidizing bacteria; AD - anaerobic digestion; VFAs - volatile fatty acids; SCFAs - short-chain fatty acids; COD - chemical oxygen demand; PCoA - principle coordinate analysis; VSS - volatile suspended solids; MAGs - metagenome assembled genomes; FBEB – flavin based electron bifurcation; RET – reverse electron transfer; PMF – proton motive force; PFL – pyruvate formate lyase; PFOR – pyruvate ferredoxin oxidoreductase; FHL – formate hydrogenlyase; EMO – membrane-bound oxidoreductase.

## Acknowledgments

This work was supported by the Zhejiang Provincial Natural Science Foundation of China under Grant No. LR22D010001 and Key R&D Program of Zhejiang (2022C03075) and Research Center for Industries of the Future (RCIF). Technical assistance from Ms. Yisong Xu in laboratory affairs and Mr. Guoqing Zhang in maintaining the in-house computing facility are greatly appreciated.

## Authors’ contributions

YW conceptualized this research and discussed it with FJ. YW conducted the wet-lab experiment, the meta-omics data analysis and visualization, and wrote the manuscript. RZ participated in the wet-lab experiment. CW, WY, TZ and FJ contributed to the proofreading of the manuscript. FJ provided funding resources for this research.

## Footnotes

1 Theoretical calculation by authors, assuming pH=7.0, 35 °C, acetate, propionate and butyrate concentration at 10 μM, 70% CH₄ and 30% CO₂ in the gaseous phase.

